# Nutrient addition on grazing lawns and selection by free-roaming mammalian herbivores in a nutrient poor savanna

**DOI:** 10.1101/2024.01.20.576488

**Authors:** Bradley Schroder, Frank Van Langevelde, Nicola-Anne Hawkins Schroder, Herbert H. T. Prins

## Abstract

Grazing lawns are important food sources in nutrient poor savannas for free-roaming mammalian herbivores. It has been hypothesized that increased grazing pressure by mammalian herbivores can create and maintain patches of lawn grass. We tested whether the application of specific nutrients, nitrogen (N), phosphorus (P) or in combination with calcitic and dolomitic lime (Ca), in a nutrient poor African savanna, would make the grass sward more nutrient rich, which would attract mammalian herbivores to graze more frequently. We investigated the grazing patterns of six species of mammalian herbivores, namely, blue wildebeest (*Connochaetes taurinus*), Burchell’s zebra (*Equus quagga burchellii*), common eland (*Taurotragus oryx*), impala (*Aepyceros melampus*), square-lipped rhinoceros (*Ceratotherium simum*) and warthog (*Phacochoerus africanus*). We show that the addition of N attracts and increases the grazing pressure for three of the herbivore species, namely, blue wildebeest, Burchell’s zebra and impala. Our findings suggest that these often abundantly present mammalian herbivores with intermediate body mass, attracted to grazing lawns by the addition of N, can maintain grazing lawns.

**Conservation implications:** Artificial fertilization with nitrogen attracts large free-roaming herbivore species to localized grazing lawns, stimulating the creation and expansion of high nutrient quality lawn grasses in nutrient poor savannas. This results in a nutrient high food source which would normally not be available in nutrient poor savannas.

## Introduction

Food quality and quantity are strongly linked to the survival and reproduction of mammalian herbivores (Gaillard et al. 2000; Uher-Koch et al. 2019). In nutrient poor savannas, the quality of grass is often below the requirements needed by mammalian herbivore survival (Demment & Van Soest 1985; Codron et al. 2007; Prins & Van Langevelde 2008). Foraging on grazing lawns in these dystrophic areas can increase the quality of their food resources (Archibald 2008; Cromsigt & Kuijper 2011; Novellie & Gaylard 2013). Grazing lawns are areas with nutritious, grazing-tolerant grass species, with a short-stature, such as Couch grass (*Cynodon dactylon*; a.k.a. Bermuda grass), created by frequent and intense grazing (Archibald et al. 2005; Grant & Scholes 2006; Cromsigt & Kuijper 2011; Arnold 2012; Hempson et al. 2014). Stoloniferous, palatable lawn grasses can only persist under continued heavy grazing (Cingolani et al. 2005; Archibald 2008; Novellie & Gaylard 2013). If grazing pressure is relaxed, tall, fast-growing grass species of low food quality will outcompete the palatable grass species (O’Connor 1994; Uher-Koch et al. 2019). Previous research has shown that grazing lawns have been established in particular localities such as nutrient hotspots, including termite mounds and old cattle kraals (Van der Waal et al. 2011a; Cromsigt & Olff 2008), recently burnt areas (Archibald 2008; Donaldson et al.2017) but also through the addition of nutrients to grazing lawns (Cromsigt & Olff 2006; Van der Waal et al. 2011a). These nutrient-enriched localities all attract mammalian herbivores and increase local grazing pressure (Cromsigt & Olff 2006; Hempson et al. 2014). The extent of patches of grazing lawns is influenced by the number of mammalian herbivores and the continued attractiveness of these patches to these animals (Archibald et al. 2005). It has been hypothesized that grazing by certain grazer species, such as squared-lipped rhinoceros (*Ceratotherium simum*) and hippopotamus (*Hippopotamus amphibius*), can result in the establishment of grazing lawns because they are non-selective bulk grazers, utilising high quantities of low-quality food (Owen-Smith 1988; Verweij et al. 2006; Waldram et al. 2008; Cromsigt & Olff 2008; Kleynhans et al. 2011). Their concentrated grazing then results in areas with short and high-quality grass.

Studies have demonstrated that free-roaming mammalian herbivores graze more frequently in nutrient rich (fertilized) grass patches as compared to nutrient poor (unfertilized) patches (Augustine et al. 2003; Cromsigt & Olff 2008; Van der Waal et al. 2011b; Mayengo et al. 2020). However, in nutrient poor savannas, it is unknown which added nutrient (nitrogen - N or phosphorus - P) attracts free-roaming mammalian herbivores to graze frequently (Schroder et al. submitted). Based on studies by Schroder et al. (submitted), fertilization of N and P resulted in higher concentrations of P in the grass leaf, thus the expectation was that the increased effect on grazing would have been larger on plots which were fertilized with P.

Fertilization experiments revealed that grasses in savannas are strongly co-limited by N and P (Penning de Vries & Djitèye 1982; Donaldson et al. 1984; Ludwig et al. 2008; Snyman 2002; Fynn & O’Connor 2005; Craine et al. 2008). For example, Van der Waal et al. (2011b) found that phosphorus and nitrogen fertilization increased leaf P and N concentrations in grasses in a semi-arid savanna, whereas Schroder et al. (submitted) found that only P concentrations increased in the leaf, not N. The response of the grass to fertilization with a specific nutrient may have consequences for the mammalian herbivores grazing from these grasses. A deficiency of N and / or P may affect the growth, health, reproduction, lactation and survival of mammalian herbivores (Mattson 1980; Murry 1995; Augustine et al. 2003; Zhong-xian et al. 2007). Limitation of either N or P requires herbivores to select patches to obtain these nutrients, especially pregnant and lactating females (Prins & Van Langevelde 2008).

In an earlier study by Schroder et al. (submitted), elevated nutrient concentrations in both soil and grass were measured after the addition of nutrients (N, P or a combination with Ca) to grazing lawns, with the result being an increase in grazing lawns. It is crucial to quantify the local grazing pressure of free-roaming mammalian herbivores, as this is important to understand and identify what the main drivers are which allow for the establishment and maintaining of patches of lawn grasses (Archibald 2008; Hempson et al. 2014; Griffith et al. 2017). In this paper, we test whether nutrient addition (N, P or a combination with Ca) results in an increase in local grazing pressure by free-roaming mammalian herbivores in a nutrient poor African savanna. Ca was added to the soil to increase the pH levels, which allows the plants to absorb quantitatively more P, which was previously fixed and unavailable (Chimdi et al. 2012; Higgins et al. 2012).

## Material and methods

The study was conducted in the Welgevonden Game Reserve (348 km^2^), situated on the Waterberg Plateau in South Africa (24°10’S; 27°45’E to 24°25’S; 27°56’E), over a period of three years (2016 – 2018). The area is classified as warm and temperate, with summer rainfall, having distinct wet and dry seasons, stretching from October to March and April to September respectively, with the mean annual rainfall recorded as 665 mm. The mean annual maximum temperature is 27.4 °C (reaching 40 °C) and the mean annual minimum temperature is14.5 °C (reaching -4 °C). Two biomes occur in the study area, namely, the Savanna Biome and Grassland Biome, but the Waterberg Mountain Bushveld vegetation type describes the area well too (Mucina & Rutherford 2006). The area has dystrophic to mesotrophic yellow-brown apedal coarse sands (Parker 2004), characterised by ferruginous soils with a low pH. Accordingly, the vegetation is dominated by nutritionally poor broadleaved savanna plant species. Mucina & Rutherford (2006) classify this area as a nutrient poor savanna ecosystem (locally termed ‘sour veld’). The area has a diverse mammalian herbivore assemblage, dominated by blue wildebeest (*Connochaetes taurinus*), Burchell’s zebra (*Equus quagga burchellii*) and square-lipped rhinoceros. Sixty-three species of free-roaming mammals have been identified in the study area, including various antelope species, mega-herbivores and predators. Welgevonden is surrounded by a game proof fence, preventing inward and outward migration.

We tested our hypothesis in a large-scale field experiment with eight sites spread throughout the reserve (Figure 1). We divided each of the sites into five plots, each minimally 300 m x 150 m. We controlled the encroachment of trees and shrubs on the various plots by mowing with a tractor and slasher to reduce the woody component competition with the grasses (Van der Waal 2010) but also to remove prior to the experiments all moribund and combustible material. The plots were treated annually between the months of January and February with either the addition of nitrogen (N at 0.25 ton/ha = 250 g/m^2^) or phosphorus (P at 0.50 ton/ha = 500 g/m^2^). We added lime in the form of calcitic lime at 1.5 tons/ha and dolomitic lime at 1.5 tons/ha: Ca at 3 tons/ha. In order to establish which plots the mammalian herbivores selected, camera traps were erected in the centre of each plot to capture the number of visits recorded per unit time of a species. Furthermore, animal droppings were recorded to estimate the number of animals located within the plots on the opposite side of the camera deployment in order to confirm the presence of these species (Figure 2).

**Figure 1.**
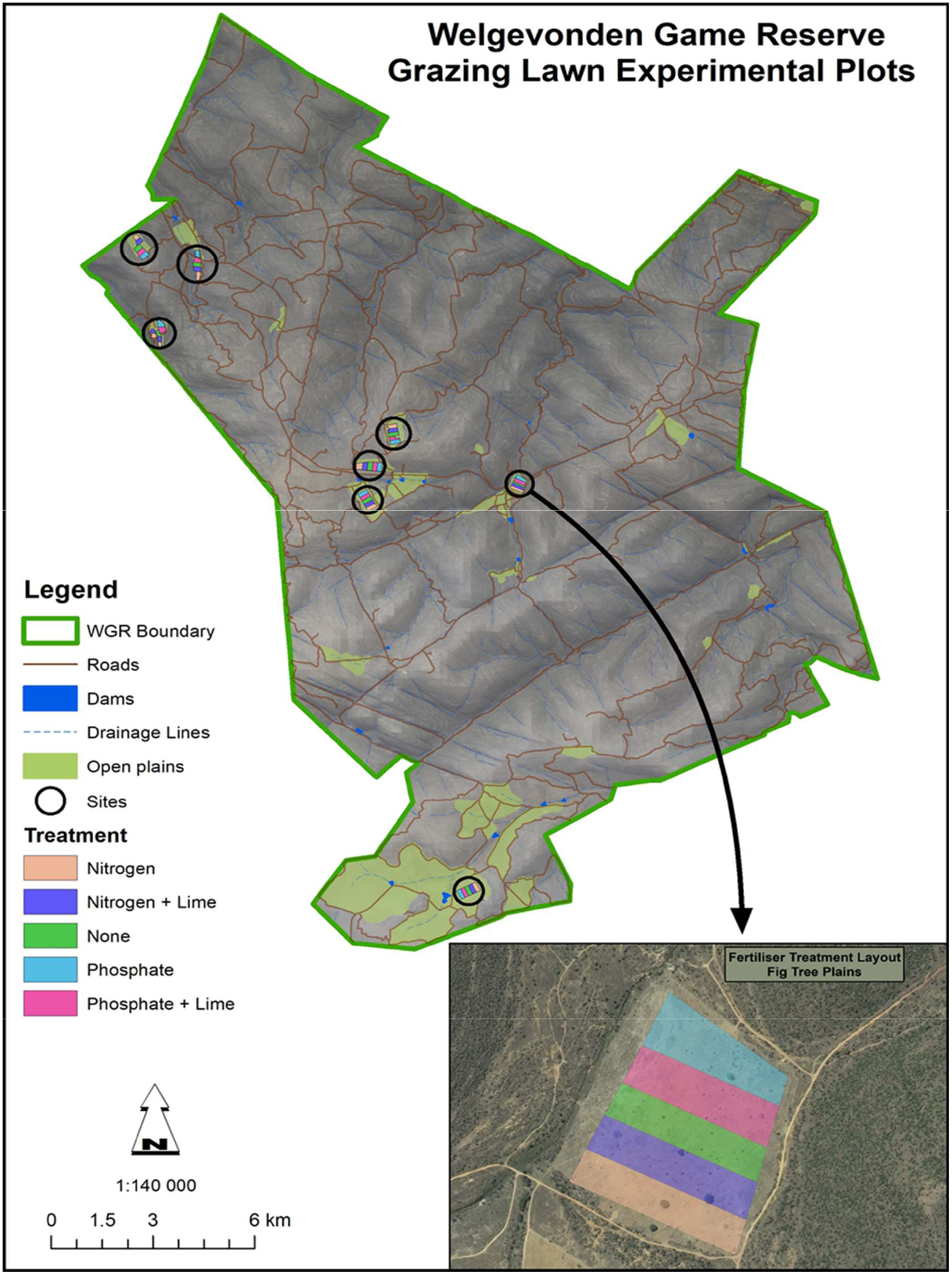
Map of the Welgevonden Game Reserve showing the grazing lawn experiment sites with the inlay of the fertiliser treatment layout at Fig Tree Plains

**Figure 2.**
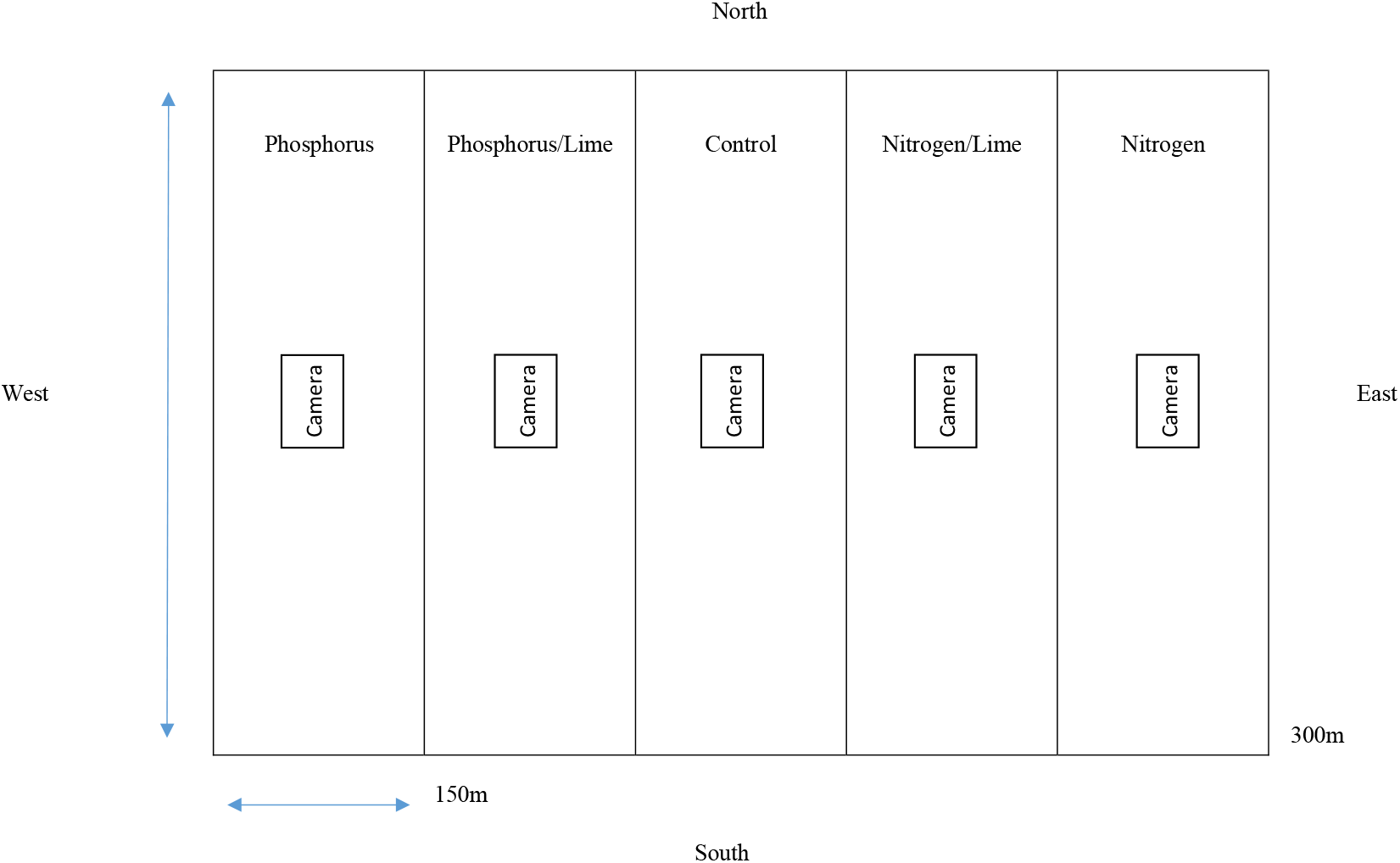
Individual site design, with five plots per site. Each plot is a representation of the different experiments undertaken with respect to the nutrients fertilized (or not). The treatments were assigned randomly. The cameras located in the middle of each plot with cameras taking photos north and south and each plot size being roughly 300m x 150m Previous studies have shown that systematic collection of faeces help to estimate population size (Plumptre & Harris 1995; Kohn et al. 1999; Barnes 2001; Vicente et al. 2004; Acevedo et al. 2007), but the method has been criticised (Ahrestani et al. 2017; Hema et al. 2017). Our design allows to test possible differences between visitation to grazing lawns, measured by camera traps and dung counts. We expect the results from the dung counts would be similar to the results established from the camera traps.

### Photographs

We used camera traps (randomized encounter model) triggered by infra-red sensors to derive relative abundance estimates, assuming that photographic capture rates, the number of visits recorded per unit time of a species, are proportional to its abundance (Carbone et al. 2001). The camera trap placements consisted of 40 Cuddeback C3-Black Flash camera traps, at roughly 1.2 meters above ground to target the majority of herbivores in the study area. We assumed that there was no difference in detection probability between the differently sized herbivores in our study (see Rowcliffe et al. 2011). The cameras were placed in the middle of each of the five plots distributed throughout the eight study sites. We sampled each plot over a two-week period, the data were downloaded after which we rotated the direction of the camera from north to south or vice versa (to obtain observations from both directions, whilst avoiding photographs being impacted by the rising or setting sun). We aligned the viewing directions of the cameras parallel to the ground, so that the horizon was in the centre of each image (Jansen et al. 2014). We deployed the cameras both during the day and the night in order to obtain equal coverage of the animal activities. Data were collected over a three-year period (2016 to 2018) during both wet and dry seasons. With each event when the camera was triggered, two sequential photographs were taken. Both images of animals were recorded as one event only. The analyses were based on these events, with 134,654 events (thus consisting of 269,308 photographs). The photographs were uploaded to Agouti, which is a platform for managing wildlife camera trapping projects. The utilisation of Agouti allowed us to organise, report and archive the images and data (Casaer et al. 2019; https://www.agouti.eu). Individual sightings of animal species were identified and enumerated to obtain a total number of each animal species identified. For each photograph we established whether an animal was grazing on the experimental plot or not. All animals of the same species were taken to be grazing if fifty per cent or more of the animals were grazing; none were considered to be grazing if less than fifty per cent of the animals were observed grazing (Mayes & Dove 2000). All photographs of objects not relevant to the study, such as humans, predators, animals with only a few recorded sightings and the like were removed prior to analysis. After analysing the data for all the mammalian herbivore species observed in the photographs, we only utilised the data for the six species that had the highest number of photos, *viz*., blue wildebeest (a grazer), Burchell’s zebra (a grazer), common eland (*Taurotragus oryx*) (a browser), impala (*Aepyceros melampus*) (a mixed feeder), square-lipped rhinoceros (a bulk grazer) and warthog (*Phacochoerus africanus*) (a grazer). The total number of individual qualifying photograph events recorded for the six species was 131,414.

### Dung counts

Concurrent with the cameras, but on the opposite side to which the cameras were facing, we collected from each plot dung samples (faecal standing crop approach), which we identified for each mammal species, enumerated and then crushed so that they would not be recorded during a following dung sampling event. This was undertaken in the same photographic range in which we placed the cameras to photograph the animals in order to obtain comparable results. We quantified one full pile of dung as a single recording (midden’s were counted as ten animals per species other than rhinoceros, which were counted as one animal per midden (Ezenwa 2004). Data were collected every two weeks over a period of two years (2017 and 2018) during both wet and dry seasons, with 62,281 individual dung samples obtained. After considering the data for all the herbivore dung samples obtained, we only quantified the data for the six herbivore species that we selected for analysis in the photographs (yielding 52,604 dung samples).

### Statistical Analysis

For each of the six grazing and browsing species (i.e., blue wildebeest, Burchell’s zebra, common eland, square-lipped rhinoceros, impala and warthog) we tested our hypothesis using generalized linear mixed models (GLMMs), using a Poisson distribution with a log link, with the number of individuals on the photos or the number of dung samples as response variable, the nutrient treatments as fixed factor, and year, period and site as random factors. The differences between the nutrient treatments were compared using Šidák multiple comparisons tests. For the number of individuals on the photos, we separated the analysis between the animals visiting the plots and those animals observed grazing to establish grazing plot preference. Analysis was performed in SPSS v. 23 (SPSS Inc., Chicago, USA).

## Results

### Photographs

The analysis of the blue wildebeest data, comprised of 21,111 photographs (covering 50,032 individual animal sightings), showed that for grazing they selected the plots treated with N (and slightly less for the plots with N in combination with Ca) (Figure 3, Table 1). The qualifying photographs for all blue wildebeest (grazing and non-grazing individuals) showed that the blue wildebeest selected the plots with N, but the differences between the treatments were small, i.e., the effect size of the N-treated plots is negligible. The data on Burchell’s zebra were comprised of 10,686 photographs (covering 14,587 individual animal sightings). The analysis of grazing Burchell’s zebra only showed that they selected the plots with N in combination with Ca (Figure 3, Table 1). For all the Burchell’s zebra, both grazing and non-grazing combined, there was no clear selection.

**Table 1.** Results of the generalized linear mixed model (GLMM) for significance between the six herbivore species (*viz*., blue wildebeest (*Connochaetes taurinus*), Burchell’s zebra (*Equus quagga burchellii*), common eland (*Taurotragus oryx*), impala (*Aepyceros melampus*), square-lipped rhinoceros (*Ceratotherium simum*) and the warthog (*Phacochoerus aethiopicus*) compared against the various treatment plots. The individual animal species are separated into observations of grazing and non-grazing. Year, period and site were the random factors, estimation methods were REML and the sample size n = 55,662 (see Figure 3 for the graphs). (df) refers to the number of degrees of freedom for the treatment levels (df1) and for those of the number of observations (df2). Model refers to the summary of the GLMM. Treatment refers to the plot fertilization with N, P, or a combination with Ca and a control. The data were collected over a three-year period, 2016 – 2018, on the Welgevonden Game Reserve, South Africa

| Blue wildebeest - grazing | F | df1 | df2 | Significance |
| --- | --- | --- | --- | --- |
| Model | 37.9 | 4 | 6.743 | <.001 |
| Treatment | 37.9 | 4 | 6.743 | <.001 |
| Blue wildebeest - non-grazing |  |  |  |  |
| Model | 3.3 | 4 | 14.358 | .011 |
| Treatment | 3.3 | 4 | 14.358 | .011 |
| Burchell's zebra - grazing | F | df1 | df2 | Significance |
| Model | 17.8 | 4 | 3.301 | <.001 |
| Treatment | 17.8 | 4 | 3.301 | <.001 |
| Burchell's zebra - non-grazing |  |  |  |  |
| Model | 5.7 | 4 | 7.375 | <.001 |
| Treatment | 5.7 | 4 | 7.375 | <.001 |
| Common eland - grazing | F | df1 | df2 | Significance |
| Model | 16.8 | 4 | 757 | <.001 |
| Treatment | 16.8 | 4 | 757 | <.001 |
| Common eland - non-grazing |  |  |  |  |
| Model | 15.6 | 4 | 2.149 | <.001 |
| Treatment | 15.6 | 4 | 2.149 | <.001 |
| Impala - grazing | F | df1 | df2 | Significance |
| Model | 16.1 | 4 | 2.428 | <.001 |
| Treatment | 16.1 | 4 | 2.428 | <.001 |
| Impala - non-grazing |  |  |  |  |
| Model | 13.4 | 4 | 7.455 | <.001 |
| Treatment | 13.4 | 4 | 7.455 | <.001 |
| Square-lipped rhinoceros - grazing | F | df1 | df2 | Significance |
| Model | 1.1 | 4 | 1.273 | .353 |
| Treatment | 1.1 | 4 | 1.273 | .353 |
| Square-lipped rhinoceros - non-grazing |  |  |  |  |
| Model | 1.0 | 4 | 1.366 | .418 |
| Treatment | 1.0 | 4 | 1.366 | .418 |
| Warthog - grazing | F | df1 | df2 | Significance |
| Model | 2.4 | 4 | 2.380 | .052 |
| Treatment | 2.4 | 4 | 2.380 | .052 |
| Warthog - Non-grazing |  |  |  |  |
| Model | 2.9 | 4 | 6.017 | .022 |
| Treatment | 2.9 | 4 | 6.017 | .022 |

**Figure 3.**
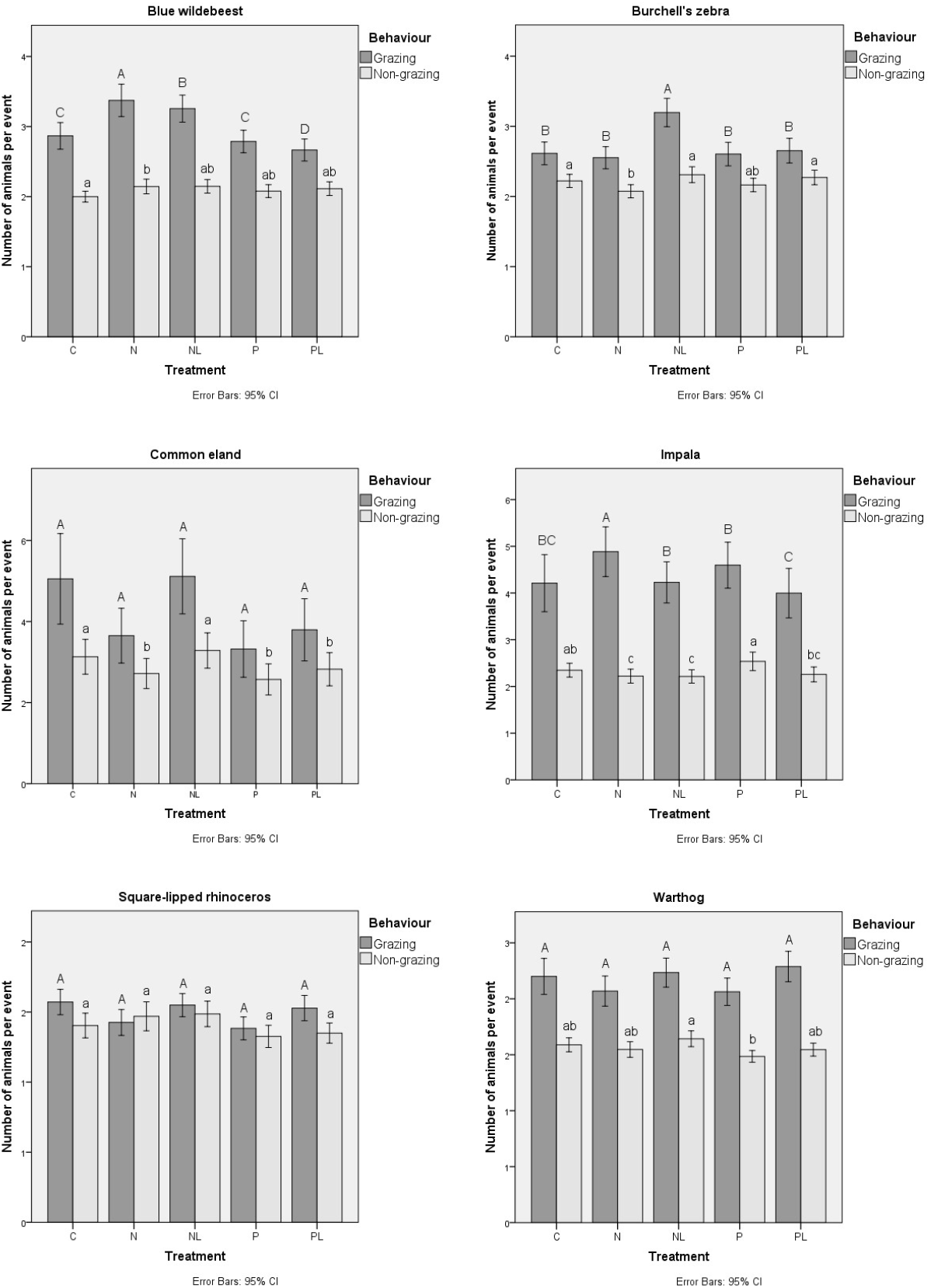
Number of animals grazing or not grazing per event for each herbivore species (*viz*., blue wildebeest (*Connochaetes taurinus*), Burchell’s zebra (*Equus quagga burchellii*), common eland (*Taurotragus oryx*), impala (*Aepyceros melampus*), square-lipped rhinoceros (*Ceratotherium simum*) and the warthog (*Phacochoerus aethiopicus*) per treatment plot, collected over a three-year period, 2016 – 2018, on the Welgevonden Game Reserve, South Africa. (C = control, N = nitrogen, NL = nitrogen and calcitic and dolomitic lime, P = phosphorus, PL = phosphorus and calcitic and dolomitic lime). Error bars represent the 95% confidence interval. Letters indicate significant differences between the treatments for grazing and non-grazing herbivores per species and per treatment based on Šidák multiple comparisons tests using a generalized linear mixed model (see Table 1 for the statistics)

The data on common eland, which encompassed 2,916 photographs (covering 9,540 individual animal sightings), did not show clear differential use of differentially treated plots either for grazing or a combination of grazing and non-grazing animals (Figure 3, Table 1). The impala data which were made up of 9,893 photographs (covering 28,031 individual animal sightings) showed that the impala grazed more in plots fertilized with N (Figure 3, Table 1) than in other plots. The non-grazing impala did not show selective use of differentially treated plots. The data on both the square-lipped rhinoceros, which were made up of 2,649 photographs (covering 3,849 individual animal sightings), and the warthog, which were made up of 8,407 photographs (covering 14,587 individual animal sightings) showed no clear differential use of differently treated plots for either grazing or a combination of grazing and non-grazing animals (Figure 3, Table 1).

#### Dung counts

The data for blue wildebeest, which comprised of 28,597 faeces samples, and those for the common eland, which consisted of 1,377 faeces samples, both showed no clear difference in the amount of dung samples between the various treatment plots and the control. The data for Burchell’s zebra, which involved of 4,033 faeces samples, showed a slightly higher number of dung samples in all the fertilized plots when compared to the control plots. The impala data, which comprised of 9,816 faeces samples, showed more dung in the plots with N, N in combination with Ca, and P. The dung data for the square-lipped rhinoceros, which comprised of 883 faeces samples, showed more dung piles in the plots fertilized with P and then N in combination with Ca. For warthog, we collected 7,898 faeces samples, higher amounts of dung were found in the plots with N in combination with Ca and P, and the amount of dung in the fertilized plots was higher than found in the control plot (Figure 4, Table 2).

**Table 2.** Results of the generalized linear mixed model (GLMM) for significance between the dung samples (faecal standing crop approach) identified for the six herbivore species (*viz*., blue wildebeest (*Connochaetes taurinus*), Burchell’s zebra (*Equus quagga burchellii*), common eland (*Taurotragus oryx*), impala (*Aepyceros melampus*), square-lipped rhinoceros (*Ceratotherium simum*) and the warthog (*Phacochoerus aethiopicus*) compared against the various treatment plots. Year, period and site were the random factors, estimation methods were REML and the sample size n = 5,112 (see Figure 4 for the graphs). (df) refers to the number of degrees of freedom for the treatment levels (df1) and for those of the number of observations (df2). Model refers to the summary of the GLMM. Treatment refers to the plot fertilization with N, P, or a combination with Ca and a control. The data were collected over a three-year period, 2016 – 2018, on the Welgevonden Game Reserve, South Africa

| Blue wildebeest | F | df1 | df2 | Significance |
| --- | --- | --- | --- | --- |
| Model | 203.2 | 4 | 1.465 | <.001 |
| Treatment | 203.2 | 4 | 1.465 | <.001 |
| Burchell's zebra | F | df1 | df2 | Significance |
| Model | 10.9 | 4 | 1.003 | <.001 |
| Treatment | 10.9 | 4 | 1.003 | <.001 |
| Common eland | F | df1 | df2 | Significance |
| Model | 2.8 | 4 | 483 | .027 |
| Treatment | 2.8 | 4 | 483 | .027 |
| Impala | F | df1 | df2 | Significance |
| Model | 41.8 | 4 | 934 | <.001 |
| Treatment | 41.8 | 4 | 934 | <.001 |
| Square-lipped rhinoceros | F | df1 | df2 | Significance |
| Model | 15.5 | 4 | 239 | <.001 |
| Treatment | 15.5 | 4 | 239 | <.001 |
| Warthog | F | df1 | df2 | Significance |
| Model | 73.7 | 4 | 958 | <.001 |
| Treatment | 73.7 | 4 | 958 | <.001 |

**Figure 4.**
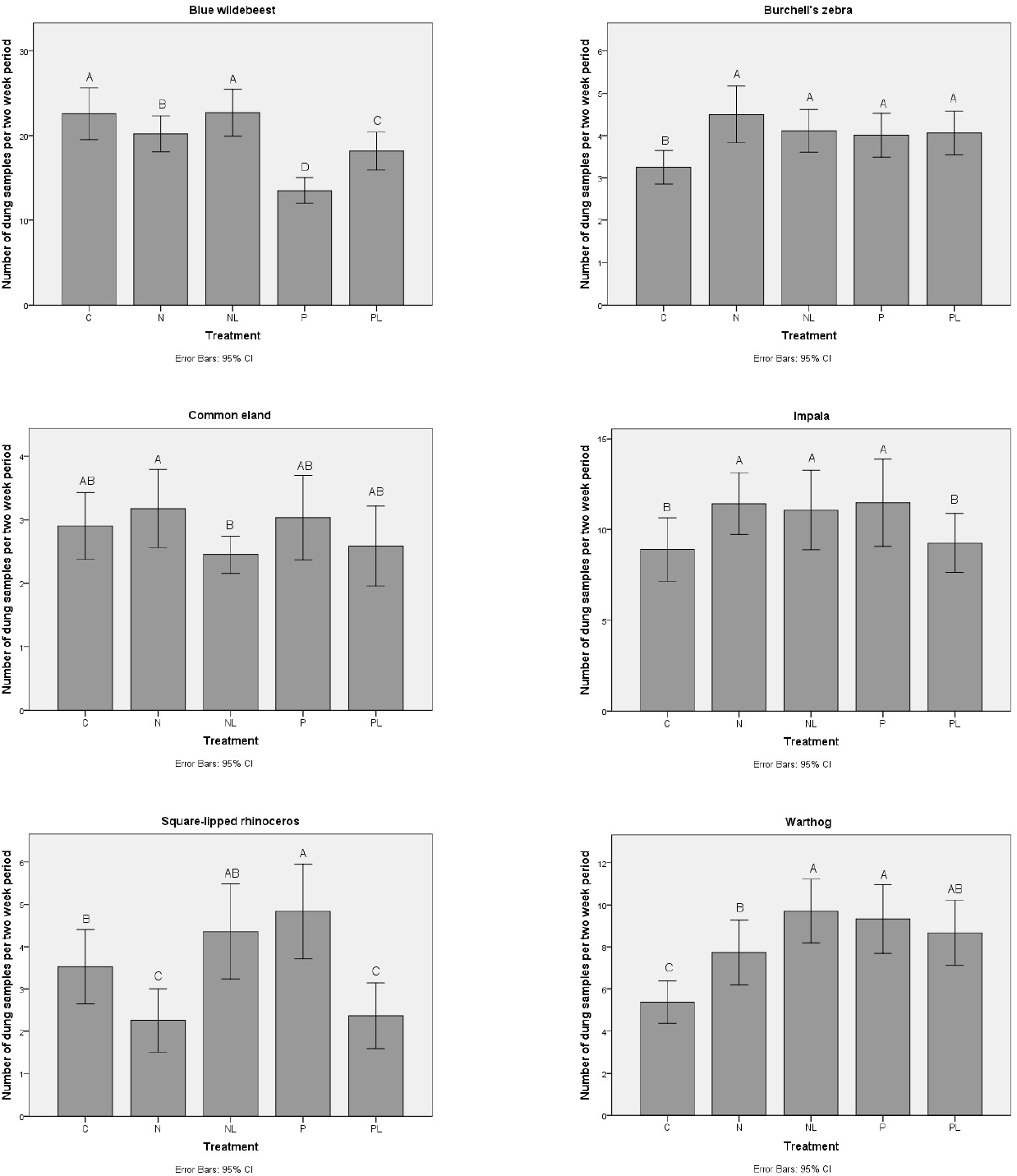
Number of dung samples observed per two-week period for each herbivore species (*viz*., blue wildebeest (*Connochaetes taurinus*), Burchell’s zebra (*Equus quagga burchellii*), common eland (*Taurotragus oryx*), impala (*Aepyceros melampus*), square-lipped rhinoceros (*Ceratotherium simum*) and the warthog (*Phacochoerus aethiopicus*) per treatment plot, collected over a two-year period, 2017 and 2018, on the Welgevonden Game Reserve, South Africa. (C = control, N = nitrogen, NL = nitrogen and calcitic and dolomitic lime, P = phosphorus, PL = phosphorus and calcitic and dolomitic lime). Error bars represent the 95% confidence interval. Letters indicate significant differences between the treatments for dung samples observed for the herbivores per species and per treatment based on Šidák multiple comparisons tests using a generalized linear mixed model (see Table 2 for the statistics)

## Discussion

In this paper, we test whether nutrient addition results in an increase in local grazing pressure by free-roaming mammalian herbivores in a nutrient poor African savanna. Therefore, we measure the presence and grazing pressure of six herbivore species on experimental plots that were fertilized by N, P or a combination with Ca in order to make the grass sward more nutrient rich to test whether grazing lawns could expand (Schroder et al.submitted). To date, research has shown that fertilization with nutrients attracts herbivores to graze, yet in nutrient poor savannas it is unknown which nutrient (nitrogen N or phosphorus P) attracts free-roaming mammalian herbivores (Schroder et al. submitted). Given the low pH in the soils of the study area, Ca was added to a number of the experiment plots, as Ca has been identified to increase the P value in grass leaves (Higgins et al. 2012). We found that after the application of the fertilizers, several herbivore species had preference for grazing in the plots with N, namely, blue wildebeest, Burchell’s zebra and impala. The addition of N combined with Ca promotes grazing by blue wildebeest and made them aggregate on these plots. Our study supports the idea that locally elevated soil nutrient levels attract grazing mammalian herbivores, especially when N is added. Furthermore, findings by Schroder et al. (submitted) and Van der Waal et al. (2011b), showed that plots fertilized with N and P, resulted in higher concentration of phosphorus (P) in grass leaves. Thus, our expectation was that the increased effect on local grazing pressure would have been larger on plots which were fertilized with P. However, our findings show that the herbivores in the study area concentrate more on grass fertilized with N and Ca, which suggests that the herbivores select forage with elevated N as it is possibly less available in the nutrient poor study area.

Camera traps were used to record the number of herbivore species visits and to quantify the behaviour of the individual species (grazing or non-grazing) on the fertilized and control plots. At the same time herbivore dung samples were recorded to estimate the number of individual animals per species present on the plots on the opposite side to the cameras. These two methods were used to establish differential use of free-roaming herbivores of sour veld vegetation (i.e., a dystrophic or nutrient poor savanna) that received different types of experimental treatment with artificial fertilizer, namely, N only, P only, a combination of N or P with Ca. We analysed more than one hundred and thirty thousand photographs of individual herbivores and more than sixty-two thousand dung samples during our study. We expected that the outcomes of the photo analysis and that of the dung samples would have yielded a similar conclusion about the use and choice of fertilized plots by the herbivores. However, this was not borne out by the results as the dung sampling did not yield an indication of differential use of fertilized vs. non-fertilized plots, while the camera trap data show that the fertilized plots are grazed more frequently. Because a priori there is no heuristic to decide which of the two methods is superior, at first sight we find ourselves in a conundrum. Indeed, Putman (1984) posits that dung sampling techniques are important for population estimation and allow for maximum possible ecological information, but Dorji et al. (2013) and Sargent (2016) criticize the use of dung sampling as a measure for local grazing pressure; they also support the notion of Silveira et al. (2003) and Morgan et al. (2019) that camera traps give a more trustworthy indication of use intensity as compared to dung sampling. Of course, we do not have the proof that camera trapping data are superior to dung sampling data, but we adhere to the notion that direct measurements of the behaviour under scrutiny (i.e., the photos of grazing animals) must be closer to the truth than indirect measurements (i.e., the dung). We thus implore for the deployment of automated picture taking equipment for determining space use by animals at fine spatial scales relative to the throughput rate of the animals under scrutiny, if one were to rely on dung pellet counting. We believe, however, that at larger scales the dung count methods are perfectly acceptable.

Our study concurs with Van der Waal et al. (2011a), in that abundantly present herbivores species are observed to frequently graze on fertilized plots. Our study shows that of the six herbivore species observed to be grazing on the lawn grass systems, the contribution per species was blue wildebeest (40%), Burchell’s zebra (20%), impala (15%), warthog (14%), square-lipped rhinoceros (7%) and common eland (4%) and shows the impact of large herbivores on these fertilized plots. Thus, our data shows that in a nutrient poor savanna, herbivorous species which we believe can induce a lawn-forming reaction of grass species have the propensity to form lawns (namely Burchell’s zebra, blue wildebeest and square-lipped rhinoceros, but not impala) can be induced to concentrate their grazing efforts after fertilizing. This concentration of grazing pressure by these herbivore species may lead to expansion of grazing lawns providing high quality food for these herbivores.

### Management implications

Artificial fertilization with nitrogen attracts large free-roaming herbivore species to localized grazing lawns, stimulating the creation and expansion of high nutrient quality lawn grasses in nutrient poor savannas. This results in a nutrient high food source which would normally not be available in nutrient poor savannas.

## Acknowledgements

The authors would like to thank the students and friends involved in helping to identify the photographs on the Agouti platform. Numerous hours were spent identifying the specific animals and their grazing behaviour. Special thanks in alphabetical order must go to Ms Coby van Dooremalen, Mr Gregory Canning, Ms Janina Harms and Ms Marit ter Bogt.

